# Mechanically induced pre-mitotic nuclear deformations promote nuclear rupture through lamin B1 depletion

**DOI:** 10.64898/2026.07.17.739104

**Authors:** Myron N.F. Hensgens, Haochun Sun, Boyd Peters, H. Martijn de Gruiter, Johan A. Slotman, Timon Idema, Hylkje J. Geertsema

## Abstract

Disassembly of the nuclear lamina is essential for successful mitosis in eukaryotic cells, but the mechanism that triggers the lamina breakdown remains unclear. Using 3D fluorescence microscopy imaging combined with 3D curvature analysis, we found that microtubule-dependent invaginations trigger lamina remodelling and lamin B1 network disruption. Monte Carlo simulations provide mechanistic insight into how those invaginations induce a mechanical discontinuity between isotropic expansion and anisotropic shrinking forces at the rim of the nucleus, which correlated with lamin B1 depletion sites and the location of cytoplasmic protein influx within the nucleus. Our findings suggest that pre-mitotic modifications of the lamina function as a mechanical trigger to facilitate nuclear rupture and are thereby a key driver of mitotic entry, complementing established biochemical pathways.

## Introduction

Structural integrity of eukaryotic nuclei is essential for genome function and maintenance and is therefore critical for cellular health. Irregularities of nuclear shape, such as nuclear ruptures and deformations, have been linked to ageing, cancer, and various diseases.^1–4^ Conversely, various vital biological processes, such as migration and cell division, impose mechanical triggers that directly influence nuclear morphology.^5^ In particular, prophase has been associated with drastic nuclear shape remodelling, followed by nuclear rupture that enables cytoplasmic proteins to enter the nucleoplasm.^6,7^ Consequently, the nucleus needs to combine mechanical stability with adaptability to support different biological processes. In this study, we aim to investigate how mechanical cues influence the reshaping and rupturing of the nucleus in prophase and thereby play a pivotal role in driving mitotic entry.

The main contributor to the structural integrity of the nucleus is a fibrous protein network called the nuclear lamina. The nuclear lamina consists of two distinct but interconnected networks that are composed of different types of lamin proteins.^8,9^ A-type lamins, comprising lamin A and its splice variant lamin C, are viscoelastic and hydrophilic, which allows them to distribute in the nucleoplasm and form a network facing the nucleoplasm.^9–12^ B-type lamins form elastic and hydrophobic networks that associate with the nuclear membrane, and are comprised of lamin B1 and B2, which are encoded by different genes.^13,14^

Current models primarily attribute nuclear lamina breakdown during the onset of mitosis to kinase-driven phosphorylation pathways, which depolymerise and redistribute lamins in the cytoplasm or endoplasmic reticulum.^15–17^ However, it remains unclear how kinases enter the nucleus, given the requirement for active transport or diffusion through holes in the nuclear envelope. Previous studies have reported that microtubule-associated nuclear indentations drive ruptures on the opposite side of the nucleus via force distributions. ^6,7^ At the same time, other studies found that lamin B network integrity is sensitive to nuclear membrane curvature, which is not drastically increased at those proposed rupture sides.^9,18–21^ The observed deep invaginations in various cell lines (e.g. neurons, HeLa cells, and BHK cells) would, however, drastically increase local curvature and, thus, could expose weaknesses in the lamin B network.^6,7,22–24^ Consequently, it currently remains unclear how mechanical processes contribute to lamin network weakening and disassembly at the onset of mitosis.

In this study, we aim to correlate the mechanical-induced changes in nuclear morphology to local lamina disruption by investigating local nuclear membrane geometry, and matching this to lamin protein levels. To do so, we employed 3D live-cell imaging and 3D confocal microscopy, combined with 3D Gaussian curvature analysis. We find that increased membrane curvature locally disrupts the lamin B1 network in non-dividing cells. During prophase, nuclear invaginations that originate from centromere-associated microtubules remodel the lamina and induce ruptures in the lamin B1 network, as we observe local influx of cytosolic proteins at those rupture sites. Interestingly, those lamin B1 ruptures are mostly found at highly curved regions of the nucleus. Monte Carlo simulations reveal that the microtubule-associated pushing forces on the elastic lamin B1 network induce a pronounced interface of anisotropic shrinking and isotropic expansion forces within the lamin B1 network, giving rise to a mechanical strain discontinuity that concentrates near the upper rim. This mechanical discontinuity within the lamin network could represent a vulnerability in the nuclear lamina, predisposing it to ruptures that enable the influx of cytoplasmic proteins and potentially initiate the known biochemical pathways. As such, our findings indicate that nuclear deformations are a key driver of lamin B1 disassembly, and thereby play a pivotal role in successful mitotic entry.

## Results

### Lamin B1 depletes at regions with increased Gaussian curvature

Previous studies have shown that nuclear shape influences 2D lamin organisation by locally reducing lamin abundance in highly curved regions, where lamin B1 was shown to be particularly curvature sensitive.^18^ We extend these studies by analysing the three-dimensional nuclear morphology and relating the nuclear shape to the local lamin density. We acquired 3D confocal microscopy images of fixed HeLa cells immunostained for lamin B1 and A/C, from which we correlated local Gaussian curvature to the average fluorescence intensity of immunostained lamin A/C and B1.

We found that lamin B1 depletes at regions with higher Gaussian curvature in 3D, while lamin A/C levels are enriched (Figure 1A). To account for variations in staining efficiency, fluorescence intensities were normalised to median values within each nucleus. Following this normalisation, lamin B1 voxel intensities at low Gaussian curvatures (0.01-0.04 μm^-2^) are normally distributed around 1.0 normalised voxel intensity (Figure 1B). At higher Gaussian curvatures (0.16-0.20 μm^-2^), lamin B1 intensity decreased substantially. In contrast, lamin A/C intensity increased modestly at intermediate Gaussian curvatures (0.04-0.10 μm^-2^), which can be attributed to lamin A/C scarring, a mechanism for nuclear envelope repair (Figure 1B).^5,12,25^ Lamin B1 intensity reduces by 15% from the lowest to the highest curvatures, while lamin A/C showed less variability, increasing by 7% at 0.07-0.10 μm^-2^ Gaussian curvature and reducing by 5% at 0.16-0.20 μm^-2^ Gaussian curvature (Figure 1C).

**Figure 1:**
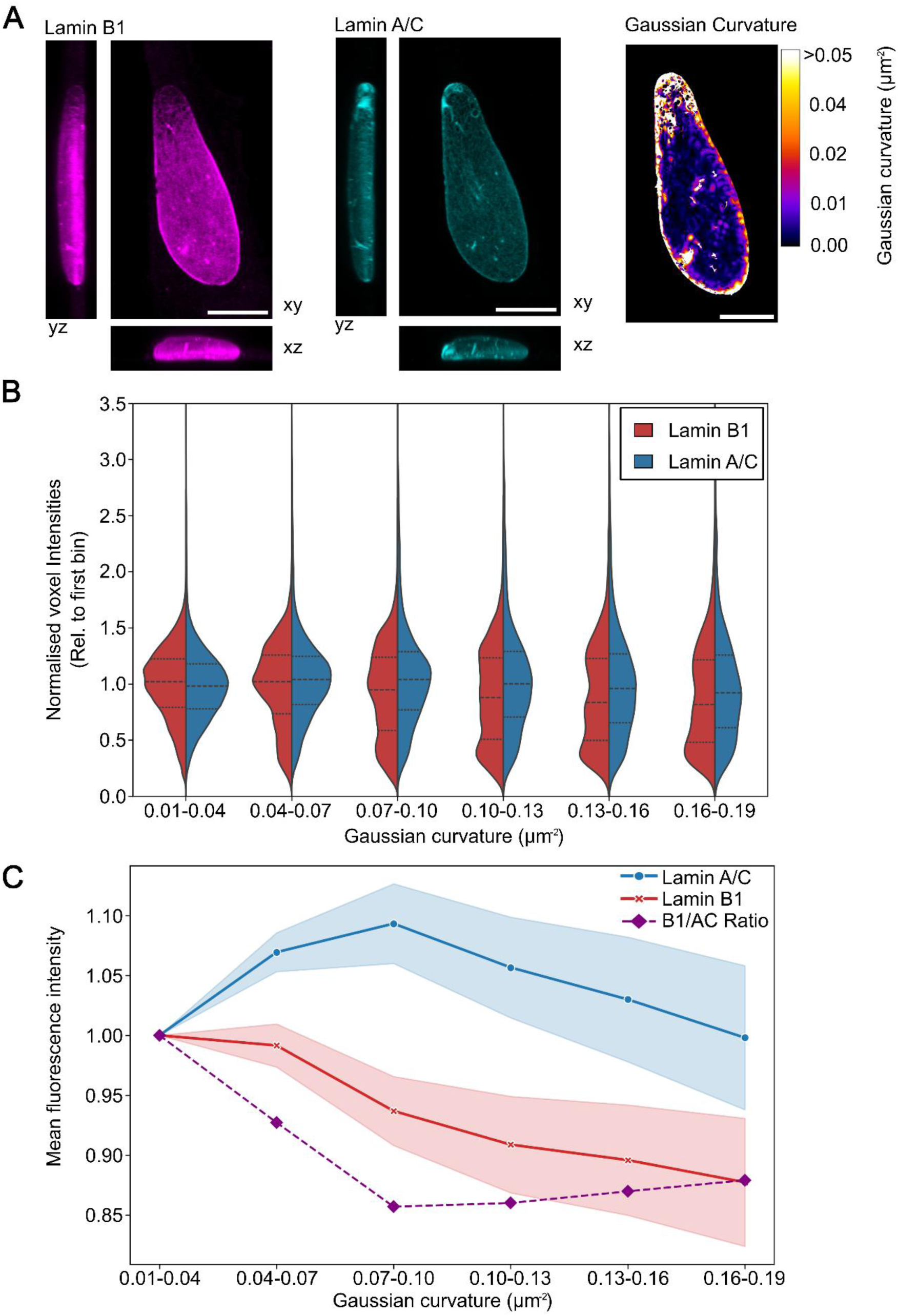
Structural integrity of lamin B1 and lamin A/C is altered at increased Gaussian curvature. **A**) Confocal image of a fixed nucleus immunostained for lamin B1, lamin A/C and the corresponding curvature heatmap. Regions with high curvature exhibit visibly reduced lamin B1 fluorescence, but not lamin A/C fluorescence. **B**) Violin plots of the voxel intensities for lamin B1 and lamin A/C, normalised to the flat nuclear regions, of six different 3D Gaussian curvature regimes. **C**) Mean fluorescent intensities of lamin A/C and lamin B1 for the different 3D Gaussian curvature regimes in blue and red, respectively, with the light colour representing the standard error of the mean. The relative depletion of lamin B1 compared to lamin A/C is shown in lilac. Scalebar in **A**, represents 10 μm. **B** and **C** were plotted with 1.2 million voxels collected from 18 different nuclei.

Additionally, ratiometric analysis demonstrated that lamin B1 is more sensitive to Gaussian curvature than lamin A/C (Figure 1C), exhibiting greater curvature-dependent depletion. Notably, since mask generation in regions fully depleted of lamin A/C and lamin B1 is not possible, holes in the lamin network with zero intensity were not included in the analysis, and the observed reductions in lamin B1 and lamin A/C could underestimate the true structural loss.

### Nuclear shape remodels at prophase due to centrosome positioning

Our results support a model in which increased Gaussian curvature imposes mechanical constraints that reduce lamin B1 density. We next asked whether such curvature-driven effects might also contribute to lamin B1 disassembly during biological processes that involve major nuclear shape remodelling, i.e. prophase. To acquire nuclear dynamics throughout the cell cycle, we performed time-lapse live-cell confocal microscopy on HeLa cells that transiently express mCherry-lamin B1. A z-stack was acquired every 10 minutes to observe nuclear morphology over extended time periods (∼12 h). Strikingly, the nuclear envelope became highly dynamic during the late prophase. The nucleus initially starts wrinkling, which is followed by the formation of a major invagination at the wrinkling site, known as the prophase nuclear envelope invagination (PNEI). Subsequently, we observe nuclear envelope breakdown, hallmarked by the disappearance of the entire nuclear envelope (Figure 2A), and later, two newly formed nuclei.

**Figure 2:**
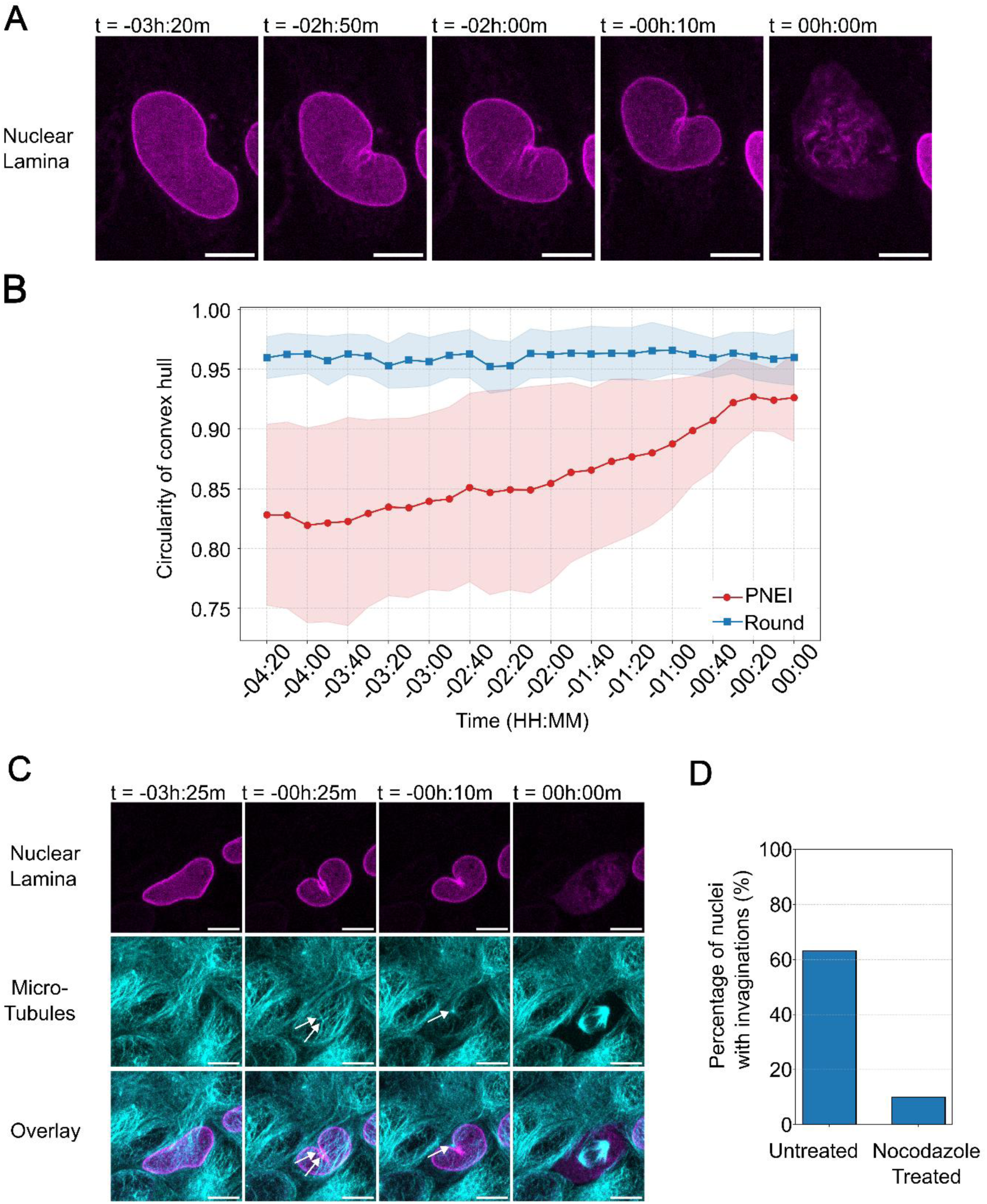
Microtubule-associated nuclear lamina remodels at late prophase. **A)** Confocal time-lapse imaging of HeLa cells that transiently express mCherry-Lamin B1 (magenta) shows nuclear lamina invaginations originating before nuclear envelope breakdown. **B)** Nuclear shape dynamics quantified as changes in the circularity of the convex hull of the nuclear lamina. Data from n=6 round and n=5 invaginated nuclei are shown over time. The shaded areas represent the standard deviation across the cell population. **C**) Confocal time-lapse imaging of HeLa cells that transiently express mCherry-Lamin B (magenta) and are labelled with SPY650-Tubulin dye (cyan), shows that centrosomes are positioned within the invagination. White arrows indicate centrosome positions over time. **D**) Percentage of nuclei with invagination before nuclear envelope breakdown of untreated (left) and nocodazole-exposed (right) cells. Total number of cells imaged without nocodazole is n=38, and the number of cells imaged with nocodazole is n=10. **A** and **C**, scale bars represent 10 µm.

We noted that initial circular nuclei exhibited PNEI formation less frequently, while initial elliptical nuclei formed a pronounced single invagination. The nuclear circularity, which is calculated as 4*_πA_*/*P*^2^, where A is the area of the convex-hull bounding polygon that bridges the invaginations, and P is the perimeter of the segmented object. We observe that the circularity increases in elliptical nuclei during prophase, as the nuclear membrane folds around the centrosomes, while initially circular nuclei remain circular throughout prophase (Figure 2B, Sup. Fig. 1). The change in shape from an elliptical to a more circular nucleus indicates significant deformation of the nucleus during prophase, supporting nuclear envelope breakdown.

Previous studies have attributed the wrinkling and invaginations to microtubules originating from the centrosome.^6,7,24^ Live-cell imaging of mCherry-lamin B-expressing HeLa cells with SPY650 tubulin dye shows that the centrosomes are located within the invagination (Figure 2C, white arrows), which suggests that centrosome-associated microtubules may drive PNEI formation and nuclear remodelling. To ensure that the nuclear deformations are indeed due to microtubule dynamics, we add nocodazole, a drug that destabilises microtubules, to our live-cell imaging experiment. Nocodazole treatment decreases the proportion of cells that exhibit invaginations from 63% in untreated cells to 10% in nocodazole-exposed cells and drastically reduces the number of successful mitotic events (Figure 2D, Sup. Fig. 1B).

### Monte Carlo simulations reveal increased force-induced curvature and strain heterogeneity in late prophase cells

Our experimental observations show that increased Gaussian curvature reduces the structural integrity of the lamin B1 network, while nuclear envelope breakdown is initiated by microtubule-associated nuclear remodelling at late prophase, which increases nuclear circularity but not Gaussian curvature. To understand how these seemingly contradictory processes can be synergistic, we aimed to clarify the mechanical dynamics of the lamin B1 network by modelling the internal force distribution within this network during prophase. We describe the lamin B1 network as a triangulated surface within the dynamically triangulated surface (DTS) framework, as implemented in FreeDTS.^26^ In contrast to standard DTS simulations of fluid membranes, only vertex displacement moves are allowed in the present model. Edge-flip (Alexander) moves are disabled to preserve mesh connectivity, thereby suppressing in-plane fluidity and rendering the membrane a solid-like elastic nuclear lamina shell.^27^

Two centrosomes are positioned at adjacent vertices on the upper surface of the nuclear shell, consistent with the observed close apposition of centrosomes at the nuclear envelope prior to mitotic entry. A constant inward force is applied to these vertices along the local inward normal, mimicking the microtubule-driven pushing of centrosomes against the nuclear envelope. The nuclear shell responds with a progressive and spatially localised deformation. At early times, the deformation remains confined near the sites of force application, resulting in a shallow indentation. As the simulation proceeds, the deformation deepens and spreads across the surface, resulting in the emergence of a ring-like region of elevated curvature in the upper part of the nucleus (Figure 3A). The corresponding Gaussian curvature heatmap quantifies intrinsic increase of curvature, which corresponds to increasing complexity and stretching of the lamina meshwork. We also show the extrinsic mean curvature to visualise the specific inward buckling caused by the modelled centrosomal invaginations (Figure 3A).

**Figure 3:**
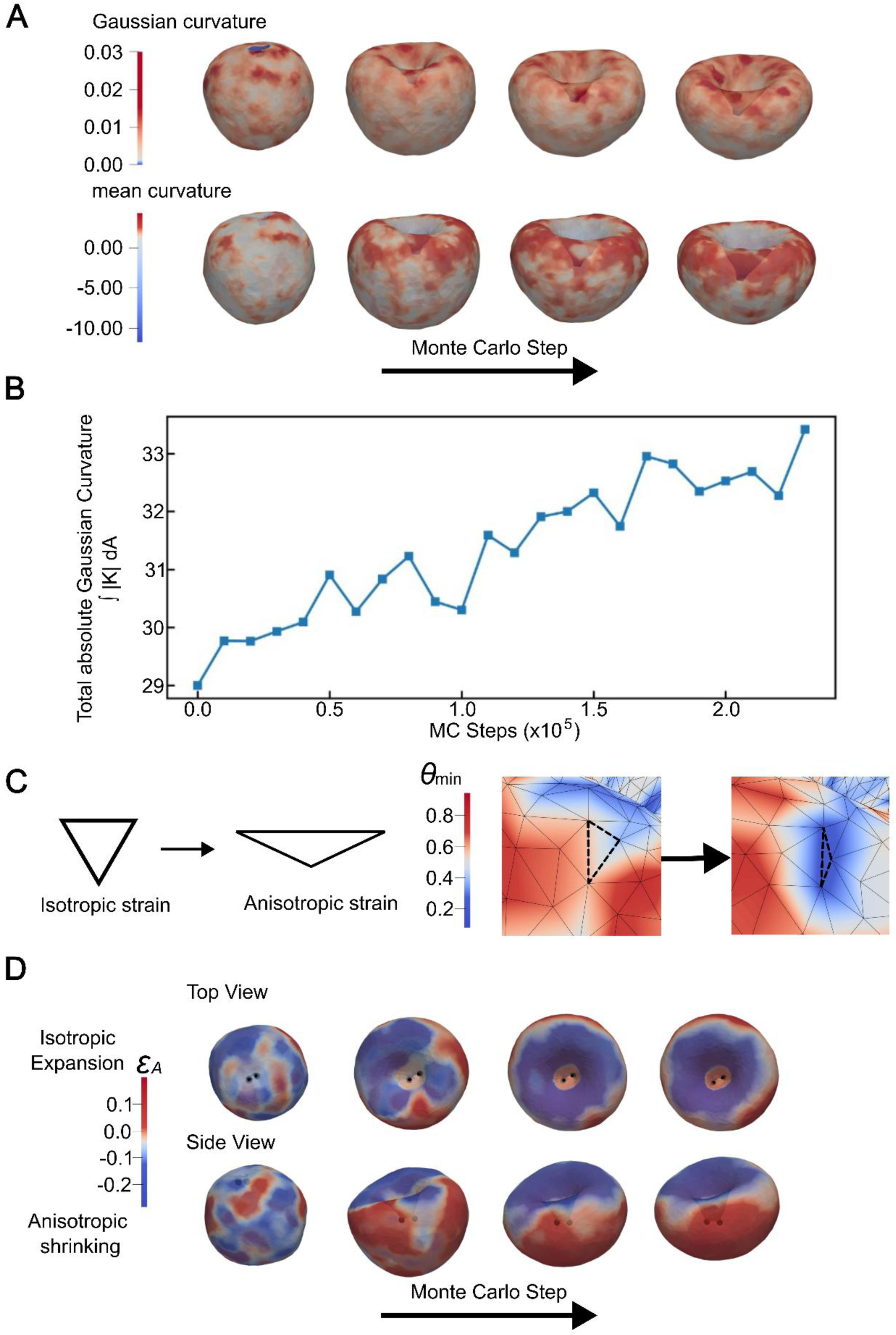
Spatiotemporal evolution of curvature and local area strain during Monte Carlo simulation. **A)** Time-lapse heatmaps of Gaussian (top) and mean (bottom) curvature. The simulation shows a progression from a spherical nucleus to an invaginated nucleus, characterised by the emergence of regions of increased curvature, especially at the rim of the centrosome-induced invagination. **B**) Total absolute Gaussian curvature over time, indicating a continuous increase in absolute Gaussian curvature. This continuous rise quantifies the accumulation of strain and tension as the mesh reorganises to accommodate the invagination. **C**) Strain anisotropy can be deduced from the shape of the triangles in the mesh. Equilateral triangles correspond to isotropic deformations, while triangles with small angles indicate large anisotropy. **D**) Local deformation (strain) in top view (top row) and side view (bottom row). Blue colours correspond to anisotropic shrinking, red to isotropic expansion, with transition between shrinking and expansion emerging and concentrating near the upper rim, where the Gaussian and mean curvatures are the highest. This border region between expansion and shrinking is where the strain gradient is largest, and we expect the nuclear lamina to break.

The temporal evolution is quantitatively captured by the total absolute Gaussian curvature. The total absolute Gaussian curvature increases continuously during deformation, reflecting the progressive accumulation of localised curvature heterogeneities (Figure 3B). The rise in localised curvature heterogeneities is driven by the transition from isotropic to anisotropic vertex strain (Figure 3C). Here, θ*_min_* denotes the smallest interior angle of each triangular face, which serves as a measure of local anisotropic strain: equilateral triangles ( θ*_min_* = 60° ) correspond to isotropic deformation, while smaller values indicate increasing anisotropy. Spatial analysis of the final configuration further reveals that regions of high curvature coincide with elevated vertex deformations, demonstrating a strong mechanical coupling between bending and stretching in the elastic nuclear model (Figure 3C).

Our Monte Carlo simulations can be used to localise the progression of shrinking and expansion (Figure 3D). At the start, the areas of shrinking and expansion are homogeneously distributed over the entire sphere, and all strains are small and isotropic. As the invagination becomes more prominent, a clear border between anisotropic shrinking and isotropic expansion becomes evident at the upper rim. The border region is where the strain gradient is largest. We propose that this microtubule-driven mechanical discontinuity at the interface between anisotropically shrunk and isotropically expanded regions may represent a vulnerability in the nuclear lamina, predisposing it to rupture under sustained centrosomal forcing.

### Regions of strain heterogeneity in lamin B1 correlate with nuclear ruptures in invaginated nuclei

The Monte Carlo simulations elucidated that PNEI progression elevated strain heterogeneity in the lamin network, with distinct regions of high strain gradient that could expose a vulnerability in the lamin B1 network. These vulnerable regions could facilitate nuclear rupture and nuclear envelope breakdown. To investigate whether locations exposed to high strain gradient correlate with nuclear ruptures, we used confocal microscopy on synchronised HeLa cells that were fixed just before nuclear envelope breakdown (NEBD), and immunostained for lamin B1 and cyclic GMP-AMP synthase (cGAS). cGAS is a cytosolic DNA-binding protein that can enter the nucleus upon nuclear rupture and bind to endogenous DNA at rupture sites (Figure 4A, Sup. Fig. 2). Interestingly, we found that most nuclei with PNEIs show reduced lamin B1 levels at the upper rim, corresponding to the region of maximum curvature and the strain interface found in the simulations. These regions of reduced lamin B1 coincide with the presence of cGAS foci (Figure 4A). Normalised fluorescence intensities of lamin B1 and cGAS along a selected line clearly show the inverse relationship between the presence of lamin B1 and cGAS, and the increase in the ratio of cGAS to lamin B1 at nuclear rupture sites (Figure 4C, D). Furthermore, the presence of cGAS sites in the nucleus was strongly reduced in spherical nuclei (11 %) compared to invaginated nuclei (78 %) (Figure 4B, Sup. Fig. 3).

**Figure 4:**
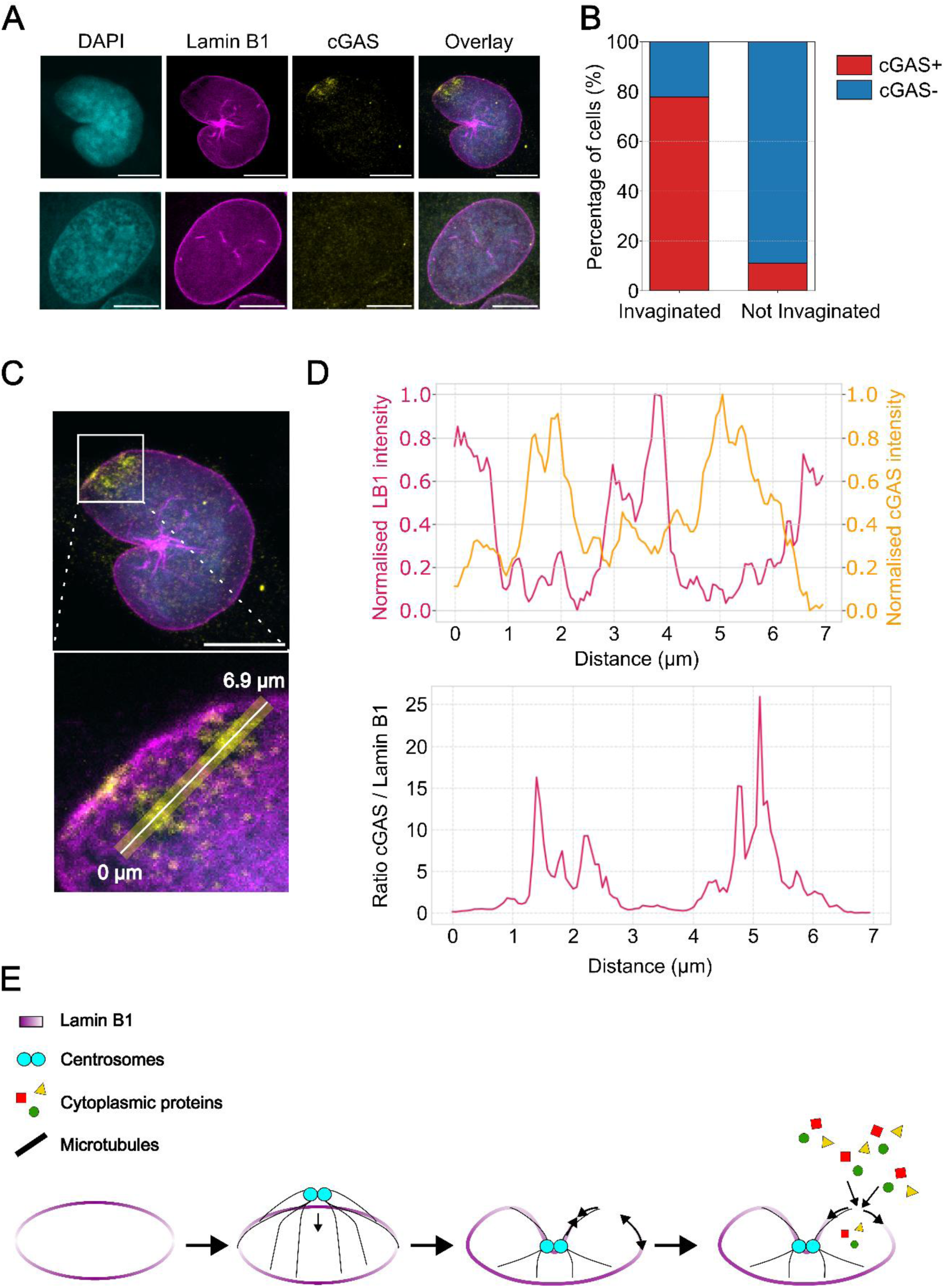
Lamin B1 depletion sites coincide with nuclear envelope ruptures. **A)** Confocal microscopy images of fixed HeLa cells, stained for DAPI (cyan), and immunostained for lamin B1 (magenta) and cGAS (yellow). The top row depicts a nucleus with an invagination, showing local depletion of lamin B1 that correlates with cGAS foci, and thus a nuclear rupture site. The bottom row shows an elliptical nucleus with a uniform distribution of lamin B1 and no cGAS foci. **B**) Bar charts of nuclear envelope ruptures, quantified by the presence of at least one cGAS focus, in invaginated versus elliptical nuclei. A significantly higher percentage of invaginated nuclei (28 of 36 cells) show nuclear ruptures than non-invaginated nuclei (4 of 36 cells). **C**) Confocal immunofluorescence image of the nucleus shown in (A), with a magnified insert showing the region of the nuclear rupture. The white line indicates the depicted line profile used for quantification in (D). **D**) The normalised fluorescence intensity for lamin B1 (magenta) and cGAS (yellow) along the line profile in (C) demonstrates an inverse relationship, as a local reduction of lamin B1 intensity directly correlates with a peak in cGAS fluorescence. The graph below shows the cGAS to lamin B1 fluorescence ratio, which is enhanced at nuclear rupture locations. **E**) Conceptual model for the onset of nuclear breakdown during mitosis. Microtubule-driven PNEI result in nuclear lamin strain, causing mechanical heterogeneity of lamin B1 at the upper nuclear rim. Consequently, the upper rim becomes vulnerable to nuclear ruptures, which in turn facilitate the influx of cytosolic proteins into the nucleus, including cGAS and kinases. Kinases could then activate further biochemical pathways that lead to full nuclear envelope breakdown. Scale bars in (**A**) and (**C**) represent 10 μm.

Taken together, our findings provide a strong indication that microtubule-driven PNEIs lead to a mechanically heterogeneous rim within the lamin B1 network that facilitates local depletion of lamin B1 and the initiation of nuclear rupture sites, as summarised in Figure 4E. Consequently, we suggest that this mechanistically driven process of nuclear rupture adds to the well-known biochemical pathways of kinase-driven nuclear envelope breakdown, thereby providing a key step in initiating mitosis.

## Discussion

In this study, we investigated the role of mechanical cues on lamina breakdown at mitotic entry. Our data reveal that the integrity of the lamin B1 network is specifically sensitive to increased local 3D Gaussian curvature and that the lamina undergoes significant remodelling throughout prophase, driven by a microtubule-dependent nuclear indentation. The formation of these indentations thereby imposes mechanical strain on the lamin B1 network, promoting lamin B1 depletion sites. Our Monte Carlo simulations of the nuclear lamina provide a mechanistic understanding of the synergy between those findings by quantifying the curvature and tension within the lamin network throughout prophase. We found that curvature mainly increases at the rim of the invagination, where we observe a clear interface between isotropic expansion and anisotropic shrinking. This region of heterogenous strain could weaken the protein network locally and expose nuclear rupture sites. Strikingly, our data indeed shows local lamin B1 depletion associated with nuclear envelope rupture sites at the rim of the invagination. Those rupture sites allow for an influx of cytoplasmic proteins, as evidenced by cGAS entering the nucleus, which could trigger the well-known phosphorylation pathways that depolymerise lamins and ultimately dissolve the nuclear membrane. Taken together, our study reveals that mechanical cues from microtubule-originated pushing on the lamin network facilitate lamina remodelling and upon sustained pushing nuclear envelope breakdown, and thereby drive successful mitotic entry.

Our study reveals an unexpected contribution of nuclear mechanics to the regulation of mitotic entry. While biochemical pathways have been studied and are recognised as the driving forces behind the nuclear envelope breakdown (NEBD), our findings suggest that mechanical remodelling of the nucleus, through microtubule-driven invagination and strain-induced lamin B1 depletion, could be a complementary biophysical pathway. This dual control mechanism could provide new insights into how cells integrate cytoskeletal forces to regulate essential cellular events, e.g. the timely progression into mitosis.

Our findings significantly contribute to our understanding of the role of PNEI formation in nuclear envelope breakdown. Interestingly, a subpopulation of the HeLa cells divided without forming the PNEI (37% of the cells), whereas the majority (63%) did (Figure 2D, Sup. Fig. 1A), suggesting that PNEI formation occurs frequently. As we could not determine axial invaginations due to insufficient axial resolution, our data presumably underestimate the frequency of PNEI existence.

Previous studies by Salina et al. and Turgay et al. have shown that the microtubule network plays an essential role in PNEI formation.^7,22^ Our data aligns with these studies, particularly the observation that nocodazole-induced microtubule destabilisation reduces the frequency of PNEIs. Hence, microtubule dynamics are critical in remodelling the nuclear envelope in prophase. In contrast to models that emphasise dynein-driven tearing of the nuclear envelope, our results highlight a strain-dependent mechanism that induces lamin B1 loss, which promotes nuclear ruptures.^22,23,28^ Further, we show that ruptures do not occur at sites opposite of the nuclear invaginations, as previously described, but rather at the nuclear rim where the interface of isotropic expansion and anisotropic shrinking is located. ^6,7^

Because PNEI is observed across diverse cell lines (BHK, HeLa and neurons), our findings suggest that PNEI acts as an upstream regulator of NEBD timing, adding a layer of mechanical control that complements established biochemical pathways.^6,7,23,24^ Further, B-type lamins are expressed in most vertebrate cell types, and lamin B1 knock-out mice are only viable for a short period after birth, illustrating that B-type lamin expression is essential for metazoan cells and that our proposed mechanisms for NEBD might illustrate a prevalent mechanical contribution to mitosis.^29^ Finally, our study has broader implications for understanding cell cycle regulation in development and disease, particularly in cancers with abnormal nuclear morphology and in laminopathies where lamina composition is altered.

## Acknowledgements

We would like to thank Aleksandra Placzek for assistance with cell culture, Tsion Abrahams for helping to optimise the transfections, and Michael Davidson for providing the mCherry-lamin B1 through Addgene.

## Funding details

This work was supported by TU Delft internal research funds and is supported by the “BaSyC – Building a Synthetic Cell” Gravitation grant (024.003.019) of the Netherlands Ministry of Education, Culture and Science (OCW) and the Netherlands Organisation for Scientific Research (NWO)

## Disclosure statement

No potential conflict of interest.

## Data availability statement

All data can be shared upon request.

## Statement on author contributions

MH and HG were involved in the conception and design of this study and wrote the original draft of this paper. MH performed experiments, analysed, and interpreted the obtained data. HS and TI designed, performed and analysed the Monte Carlo simulations and wrote the corresponding part of this manuscript. BP conceptualised and designed the 3D curvature extraction pipeline. MdG assisted with live cell imaging. JS, TI, HG supervised the research. All authors critically revised the intellectual content, approved the final version of this manuscript and agreed to be accountable for all aspects of this work.

## Materials and Methods

### Tissue culture

HeLa cells (DSMZ, ACC 57) were cultured in Dulbecco’s Modified Eagle Medium (DMEM) (Gibco), supplemented with 10% (v/v) fetal bovine serum (FBS) (Aventor), 1% (v/v) penicillin-streptomycin (Gibco), and 1% (v/v) 100x GlutaMAX (Gibco). Cells were cultured in 10 cm Petri dishes at 37 °C in a humidified incubator with 10% CO2. Cells were passaged upon reaching approximately 90% confluency. For live-cell imaging, cells were seeded onto glass coverslips #1.5 (Marienfeld) one day prior to transfection. For immunostaining experiments, cells were seeded onto coverslips one day before fixation.

### Transfection

The Plasmid encoding mCherry-LaminB1-10 was a kind gift from Michael Davidson (Addgene plasmid # 55069 ; http://n2t.net/addgene:55069 ; RRID:Addgene_55069).

Cells seeded on coverslips #1.5 (Marienfeld) one day prior to transfection were transfected with Lipofectamine 2000 (Invitrogen, 11668-030). 1 μL lipofectamine 2000 was diluted in 100 μL serum-free opti-MEM and incubated for 5 minutes at room temperature. Subsequently, 0.5 μg of plasmid was added to the mixture and incubated for 20 minutes to allow liposome-plasmid complex formation. Meanwhile, the culture medium was refreshed with fresh DMEM. After 20 minutes, the transfection medium was added dropwise.

### *Immunostaining* & DAPI staining

Cells were washed with phosphate-buffered saline (PBS) (Gibco) and fixed with −20 °C ethanol absolute for 10 minutes. Fixed cells were then washed three times with PBS and permeabilised using permeabilization buffer (PBS supplemented with 0.2% (v/v) Triton-X100). After permeabilization, cells were blocked with blocking buffer (5% (v/v) Goat serum in PBS) for 1 hour at room temperature.

Primary antibodies were diluted in staining buffer (1% (w/v) Bovine serum albumin (BSA)) and incubated with the cells for 1 hour at room temperature. Coverslips were then washed three times with PBS before incubation with secondary antibodies diluted in staining buffer, also for 1 hour at room temperature. After the secondary antibody staining, cells were washed three times with PBS and post-fixated with 2% (v/v) paraformaldehyde for 10 minutes at room temperature. Post-fixed cells were washed with three times with PBS and stained with 300 nM 4’, 6-diamidino-2-phenylindole (DAPI) for 2,5 minutes. Subsequently, the DAPI-stained cells were washed three times with PBS and mounted in Prolong^TM^ Glass Antifade Mountant (Invitrogen, P36980) and cured for minimal of 3 days before they were imaged. Cells used for curvature extractions were not stained with DAPI and not mounted in mounting medium.

**Table 1:** Primary antibodies used in the research.

| Table 1: Primary antibodies used in the research |  |  |  |
| --- | --- | --- | --- |
| Antigen specificity | Animal | Dilution ratio | Product |
| Lamin A/C | Mouse | 1:50 | Santa cruz, sc-376248 |
| Lamin B1 | Rabbit | 1:200 | Proteintech, 12987-1-AP |
| Lamin B1 | Mouse | 1:200 | Proteintech, 66095-1-IG |
| cGAS | Rabbit | 1:400 | Proteintech, 26416-1-AP |

**Table 2:** Secondary antibodies used in the research.

| Table 2: Secondary antibodies used in the research |  |  |  |
| --- | --- | --- | --- |
| Antigen specificity | Animal | Dilution ratio | Product |
| Anti-Rabbit Cy3 | Goat | 1:400 | Proteintech, SA00009-2 |
| Anti-Mouse Alexa Fluo 488 | Goat | 1:400 | Abcam, AB150113 |

### Fixed and Live cell imaging

Live-cell imaging was performed using a Leica SP8 confocal microscope with a 63x immersion oil objective (1.4 NA, HC PL APO CS2) and a stage-top incubator (Okolab). Prior to imaging, the microscope stage was equilibrated to 37 °C and 5% CO_2_ to maintain physiological conditions.

HeLa cells were imaged in phenol red-free DMEM (Gibco), supplemented with 1:2000 dilution SPY650-Tubulin dye (Spirochrome, SC503) to visualise microtubules. Cells exhibiting efficient transfection were selected for long-term imaging (∼12 hours). Z-stacks were acquired with 0.6 μm intervals for 5-minute time intervals and z-stacks with 0.45 μm intervals for 10-minute time intervals, depending on the desired spatiotemporal resolution. Fixed and immunostained cells used for curvature-intensity extraction were imaged with a Leica SP8 confocal microscope using a 63x immersion oil objective (1.4 NA, HC PL APO CS2). Imaging was performed with a pixel size of 80 nm in the XY plane and Z-stack intervals of 160 nm, optimised for subsequent curvature extraction.

### Curvature and fluorescent intensity extraction

Our novel image-processing pipeline begins with 3D confocal images of fixed cells immunostained for lamin B1 and lamin A/C. Nuclei were manually segmented and their labels were expanded without overlap. Fluorescence intensities were normalised to the full intensity range of the uint8 image data (0–255).

To obtain isotropic resolution, images were convolved in the xy-dimensions with a Gaussian kernel to match the axial resolution, determined by the longest excitation wavelength (561 nm). The required Gaussian standard deviation was estimated by simulating the point spread function under imaging conditions using the Gibson-Lanni.^30^ Gaussian functions were fitted to both lateral and axial PSF line profiles using nonlinear least-squares fitting, and the resulting standard deviations were used to determine the standard deviations for blurring. Following blurring, the images were down-sampled laterally by intensity summation, resulting in isotropic voxel dimensions. A voxel-wise maximum-intensity projection across both channels was then computed, preserving the highest signal at each voxel to best represent the local nuclear lamina structure.

A rough nuclear lamina mask was generated using an unsharp-masking approach followed by local thresholding, adapted from Righolt et al. (2011), with the modification that blurring was performed isotropically.^31^ Specifically, the Gaussian blur standard deviation was set to 1.0 µm, α to 0.85, and the intensity cutoff factor (I_c,factor_) to 0.4. The cutoff intensity (I_c_) was defined as the maximum intensity of the blurred mage multiplied by I_c,factor_. β was computed as I_c_*(1 – α).

Subsequently, a 3D optimal steerable surface filter was applied to the image with a scale of 0.35 µm. The filter response was subjected to non-maximum suppression (NMS).^32^ The resulting NMS response was normalized based on the local response mean and variance estimated via Gaussian smoothing (σ = 0.7 µm). Voxels exceeding a threshold (z > 1.0) within the rough mask were retained. Finally, only the largest connected component was preserved, yielding the final nuclear lamina mask.

For the curvature analysis, the final mask was smoothed using a Gaussian kernel (σ = 1 voxel). Unsigned principal curvatures were then computed using a Knutsson mapping-based approach, which is well suited for shell-like geometries and enables robust estimation of curvature directly from grayscale images.^33^ Specifically, σ_g_ was set to 1 voxel (160 nm), σ_t_ was set 3.5 voxels (556 nm), and σ_k_ was set to 1 voxel (160 nm). The local absolute Gaussian curvature was calculated as the product of the two unsigned principal curvatures.

To relate the curvature to protein distribution, local fluorescence intensity was quantified by computing the mean signal within a spherical neighbourhood (radius 0.6 µm) centered at each voxel within the nuclear mask.

Finally, voxels were binned based on their absolute Gaussian curvature. For each nucleus, the mean intensity of the lowest Gaussian curvature bin was computed and used as normalization factor. All voxel intensities of a nucleus were divided by this value to enable comparison across nuclei.

### Simulation methods

The nuclear envelope is modelled as a triangulated surface within the dynamically triangulated surface (DTS) framework, as implemented in FreeDTS.^26^ In contrast to standard DTS simulations of fluid membranes, we only allow vertex displacement moves. Edge-flip (Alexander) moves are disabled in order to preserve mesh connectivity, which effectively suppresses in-plane fluidity and renders the membrane a solid-like elastic shell.

All quantities are expressed in reduced units. The unit of length is set by the characteristic mesh spacing *_l_*_0_, and the unit of energy is given by the thermal energy *_kBT_*. Accordingly, areas scale as *l_0_*^2^, while curvatures scale as 1/*_l_*_0_. To maintain mesh regularity, a hard constraint is imposed on the edge length *_l_*, which is restricted to the range *_l_* ∈ [1, √5] *_l_*_0_. Trial moves that result in edge lengths outside this range are rejected outright.

The nuclear shell is represented as a triangulated surface, where local geometric properties are evaluated at the vertex level. For each vertex *_i_*, the two principal curvatures *_c_^i^*_1_ and *c^i^*_2_ are computed using a discrete shape-operator approach. The mean (H) and Gaussian (K) curvatures are defined as

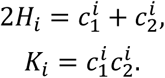

The total energy of the system is given by

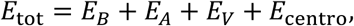

where *E_B_* is the bending energy, *E_A_* and *E_V_* impose constraints on the total area and enclosed volume, respectively, and *E*_centro_ accounts for centrosome–nucleus interactions.

The bending energy is described by a discretized Helfrich Hamiltonian^34^:

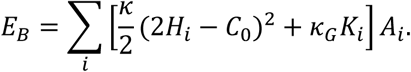

Here, *κ* and *κ_G_* are the bending rigidity and Gaussian modulus, respectively, *C*_0_ is the spontaneous curvature, and *A_i_* denotes the area associated with vertex *i*.

To regulate the total surface area, we introduce a harmonic penalty on deviations from the target area:

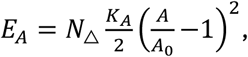

where *A* is the total area, *A*_0_ is the target area, and *K_A_* is the area compressibility modulus, and N_△_is the total number of triangular faces Volume conservation is enforced through a quadratic penalty:

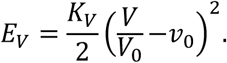

Here, *V* is the enclosed volume, *V*_0_ = *A*_0_^3/2^/(6√*π*) is the volume of a sphere with surface area *A*0, *v*_0_ ∈ (0,1] is the target reduced volume, and *K_V_* is the volume coupling constant. The effect of centrosomes on the nuclear envelope is incorporated through a curvature-dependent modification of the local elastic energy at centrosome-associated shell regions:

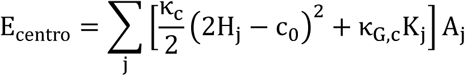

where the sum runs over centrosome-associated vertices *j*, and *κ_c_*, *κ_G_*_,*c*_, and *c*_0_ denote the local bending rigidity, Gaussian modulus, and spontaneous curvature at centrosome sites, respectively.

To further capture the mechanical action of centrosomes — such as microtubule-driven pushing or pulling on the nuclear envelope — a constant inward force is applied to each centrosome-associated vertex *j*:

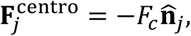

where *F_c_* is the force magnitude and **n̂***_j_* is the outward unit normal at vertex *j* . For a trial displacement Δ**x***_j_*, this force contributes an effective energy bias

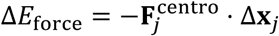

to the Metropolis acceptance criterion. Although **n̂***_j_*varies with vertex position, treating the per-move work as an energy bias is a standard approximation in membrane Monte Carlo simulations and does not introduce a conservative potential into *E*_tot_.

The system evolves via a Monte Carlo (MC) scheme using the Metropolis algorithm. At each MC step, a trial vertex displacement is proposed and accepted with probability

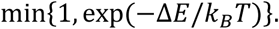

where Δ*E* = Δ*E*_tot_ + Δ*E*_force_ , and Δ*E*_tot_ is the change in total potential energy due to the displacement. We set *k_B_T* = 1 throughout, so all energies are reported in reduced units. All geometric quantities and energy contributions are updated locally after each accepted move. Simulations are run until steady-state configurations are reached.

The nuclear shell is modelled with bending rigidity *κ* = 100 *k_B_T*, Gaussian modulus *κ_G_* = 0, and vanishing spontaneous curvature *C*_0_ = 0. Mesh regularity is maintained by a harmonic spring of stiffness *k_s_* = 10 *k_B_Tl*_0_^−2^ applied to every edge with natural length *l*_eq_ = (1 + √5)/2 ≈ 1.62 *l*_0_. The total surface area is regulated by a per-triangle harmonic constraint with stiffness *k_A_* = 10 *k_B_T* and target area *A*_0_ = (1 + 2*γ*)*N*_△_√3/4 *l*_0_^2^, where *γ* = 0.81. Volume conservation is enforced by a harmonic penalty with *K_V_* = 50000 *k_B_T*(=500*κ*) and target reduced volume *v*_0_ = 1, corresponding to a spherical reference shape.

Two centrosomes are included in the simulation, each pinned to a single vertex on the nuclear shell. Each centrosome vertex is assigned bending rigidity *κ*_c_ = 20 *k_B_T*, Gaussian modulus *κ_G_*_,*c*_ = 0, and isotropic spontaneous curvature *c*_0_ = −1 *l*^−1^. A constant inward force *F_c_* = 10000 *k_B_Tl*_0_^−1^(=100*κl*_0_^−1^) is applied along the inward normal at each centrosome vertex. All simulations are run for *N* = 10^6^MC steps with a maximum vertex displacement of *δx* = 0.05 *l*_0_.

## Notes

### Competing Interest Statement

The authors have declared no competing interest.

