## Supplementary material for "Mechanically induced pre-mitotic nuclear deformations promote nuclear rupture through lamin B1 depletion": Supplemenatry Information, Figures S1-S3

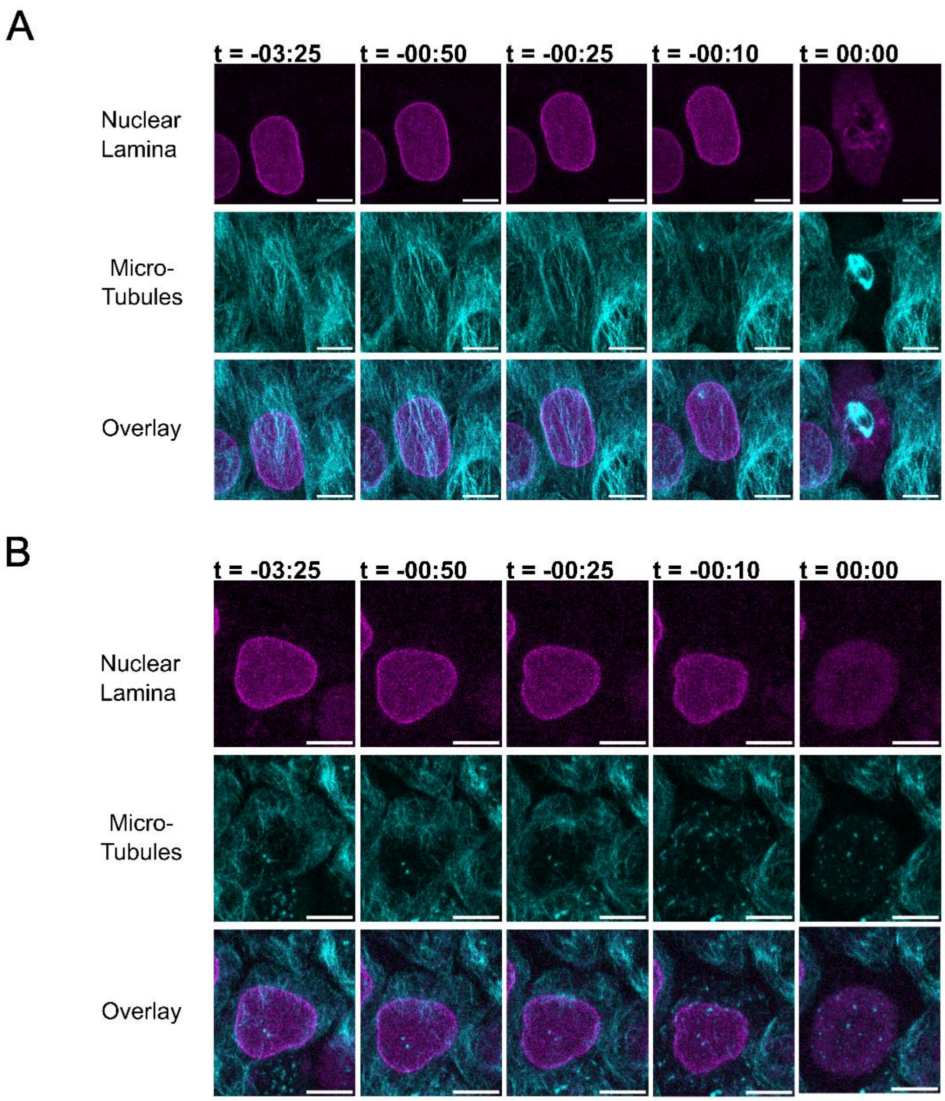


Supplementary Figure 1: Divisions without prophase nuclear envelope invaginations
**A**) Division of a HeLa cell characterised by the absence of a prophase nuclear envelope invagination without the addition of nocodazole. **B**) Nuclear envelope breakdown with the addition of nocodazole. Scale bars represent 10 µm.

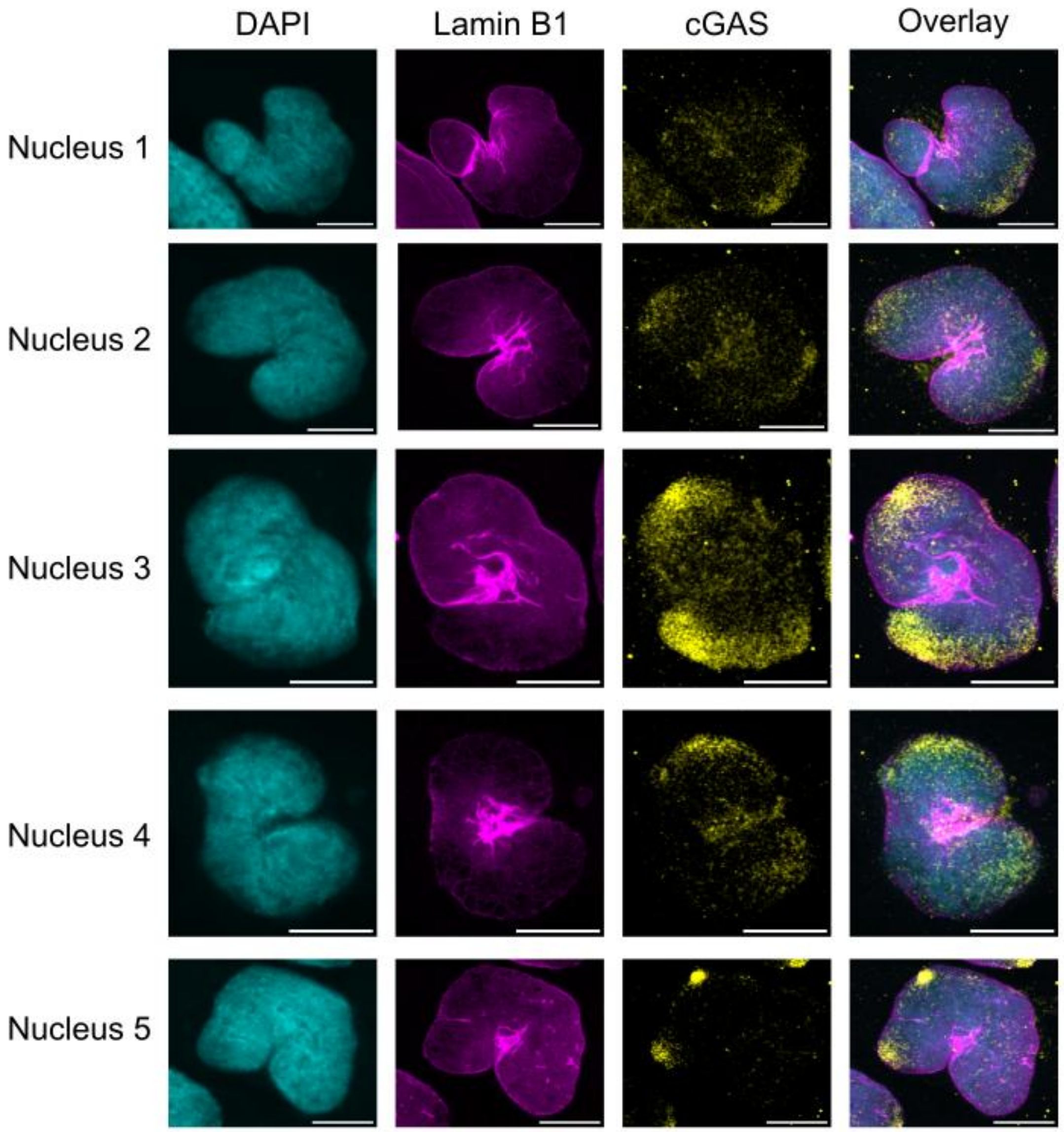


Supplementary Figure 2: Prophase nuclear envelope invaginations stained with cGas and lamin B1

Cells with prophase nuclear envelope invaginations frequently contain cGAS, a marker of nuclear envelope rupture. Scale bars represent 10 μm.

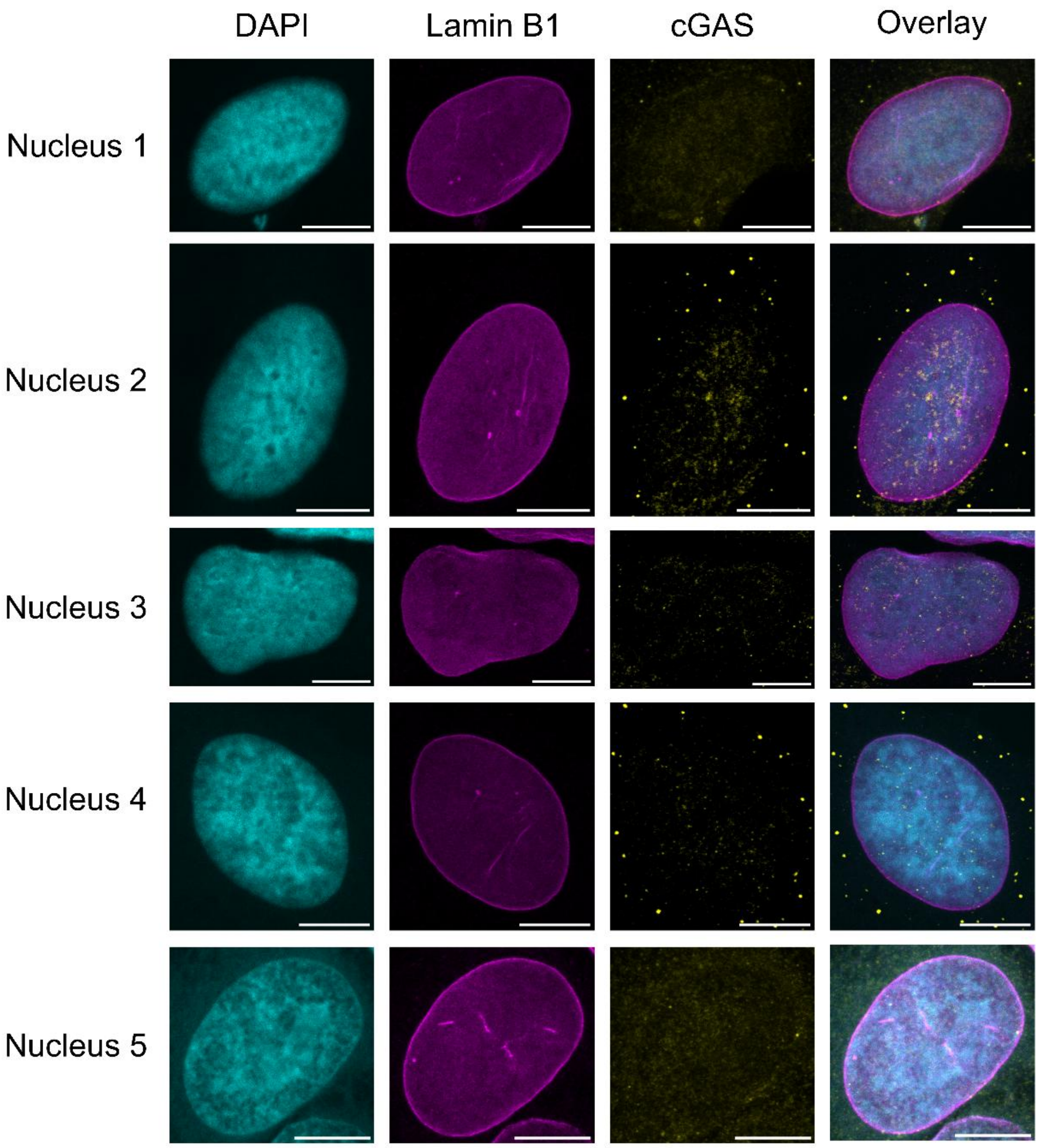


Supplementary Figure 3: Prophase nuclear envelope invaginations stained
Cells without prophase nuclear envelope invaginations do not contain cGAS foci, and therefore, do not show the characteristics of a nuclear envelope rupture. Scale bars represent 10 μm.
